# Colonizations drive host shifts, diversification of preferences and expansion of herbivore diet breadth

**DOI:** 10.1101/2020.03.31.017830

**Authors:** Michael C Singer, Camille Parmesan

## Abstract

Dynamics of herbivorous insect diet breadth are important in generation of novel pests, biological control of weeds and as indicators of global change impacts. But what forces and events drive these dynamics? Here we present evidence for a novel scenario: that specialization increases in persistent populations, but that, at the species level, this trend is countered by effects of colonizations. Colonizations cause host shifts, which are followed by non-adaptive evolutionary expansions of diet breadth, adding transitory hosts during adaptation to the principal novel host.

We base this thesis on long-term study of 15 independently-evolving populations of Edith’s Checkerspot butterfly, eight of which used fewer host genera in recent censuses than in the 1980’s, while none used more - a significant increase in specializaton. At the same time, two extintion/recolonization events were followed by temporary expansions of diet breadth. Behavioural experiments showed that these expansions were driven by within-population diversification of individual oviposition preferences. These results may explain an old puzzle: a significant negative association between population-level diet breadth and mtDNA diversity. Populations with fewer mtDNA haplotypes had broader diets, suggesting that diet breadth increases in younger, recently-colonized populations.

A recent global meta-analysis of butterfly diets, using biogeographic data, explains latitudinal patterns of diet breadth by showing that poleward range expansions have caused reduced specialization. This implies broad applicability of our results, which provide a plausible mechanism for the latitudinal trends: colonizations at expanding range margins would increase population-level diet breadths, while population persistence in range interiors would facilitate increasing specialization.

---

Evolution along a specialist-generalist axis has both practical and conceptual significance^1-3^, and herbivorous insects have figured prominently in studies of diet breadth evolution. Phylogenetic analyses have tested the plausible hypothesis that specialists are derived from generalists more frequently than evolution in the opposite direction. The hypothesis was not well supported^4,5^; evidently diet breadth evolves readily in either direction. The idea that this bidirectional evolvability causes oscillations between specialization and generalization, and that these oscillations have acted as important drivers of insect speciation and biodiversity first emerged from analyses of the butterfly family Nymphalidae^5,6^. This idea has stimulated lively and apparently unresolved debate^7-11^.

Hardy^9^ asks “does experimental adaptation of a plant-eating insect population to a novel host result in host-use generalism, and improve the odds of evolving additional new host associations?” Braga et al.^12^ use an experiment “in silico” to answer this question in the affirmative. Here we answer it “in vivo,” applying a combination of observation and experiment to a single butterfly species, Edith’s checkerspot *(Euphydryas editha)*, and showing that habitat colonizations were followed by diversification of individual host preferences and increases of population-level diet breadth.

Meta-analysis produces the generalization that mean diet breadth of insects increases with latitude^13,14^. As in other taxa^14^, temperate zone insects tend to be less specialized than those from equatorial climes. A possible reason is that generalists have been better colonists, quicker to extend their ranges polewards as glaciations receded. Using a global analysis of butterfly diets, distributions, and range dynamics, Lancaster^14^ builds a case for the opposite cause and effect: that the process of range expansion itself has generated the broader diets observed at higher latitudes. Here, we show that, in our study insects, the fine-scale mechanics of diet-breadth dynamics generate an expectation of the global patterns that Lancaster documents. As range shifts caused by climate warming increase in both pace and prevalence^15^, the combination of our behavioural study with Lancaster’s global meta-analysis will help understand changes of niche breadth that occur within these shifting ranges.

## Study system

Edith’s checkerspot butterfly (*Euphydryas editha; Nymphalidae, Melitaeinae*), uses different host genera in a geographic mosaic across its range^16-18^. Adults lay eggs in clutches on hosts in the Orobanchaceae (*Pedicularis, Castilleja*) and Plantaginaceae (*Collinsia, Plantago, Penstemon, Veronica, Mimulus, Antirrhinum*). When the proportion of *E. editha* eggs laid on each host was ascertained by census at each of 57 sites, 43 populations were recorded as monophagous, with the remainder using two to four host genera^17^. These populations showed strong isolation by distance but no isolation by host, so they did not comprise a set of host-associated cryptic species^19^.

## Relationship between population-level diets and host preferences of individual females

Because female *E. editha* behave naturally in staged encounters with potential hosts, an experimenter can assess oviposition preferences by arranging a sequence of such encounters^20-25^. Use of this behavioural assay has shown that, in populations of *E. editha* that used more than one host, this diversity of diet could be achieved either by weakness of oviposition preference (allowing butterflies to accept hosts that they did not prefer), and/or by diversity of preference rank within the population^20-22^. Diversity of rank was an important source of diet variation within two populations where diet was rapidly-evolving, while weakness of preference was the principal mechanism in 6 populations that were not currently indulging in bouts of diet evolution^22^.

## Variation within and among host populations and species: can a host shift be intraspecific?

Preference tests performed on *E. editha* and two closely-related Melitaeine species, *Melitaea cinxia* and *Euphydryas aurinia*, examined butterfly responses to variation among host individuals, populations and genera. From the perspectives of all three butterfly species, variation of acceptability among host individuals or conspecific populations can be equivalent in magnitude to variation among host genera^23-25^ (See Glossary for definitions of “preference” and “acceptability” and supplemental Text 1 for experiments). Because variation among host populations is so important to Melitaeines, it may often be the case that a colonizing female is effectively undertaking a host shift even if the host she uses after migrating is the species on which she developed at her site of origin. Host shifts may be much more frequent from the butterflies’ perspective than they appear to a human observer who classifies hosts by species.

## Results

### Long-term observations

Distributions of *E. editha* eggs and young (pre-dispersal) larvae on their hosts were recorded in the 1970’s/1980’s and again more recently in 15 populations/metapopulations distributed across California. Figure 1 shows the locations and diets of 14 of these populations and Figure 2 shows time-trends of diet breadth across decades at all 15 sites. Supplemental Table 1 complements Figure 2, giving recorded diet breadths at first and last census in each site. As the Figure shows, seven populations had the same diet breadth in the most recent census as in the 1980’s, while eight had narrower diets. None had broader diets. A two-tailed binomial test rejects the hypothesis that diet breadth was equally likely to have expanded or contracted (p = 0.008). Within our set of study populations, there has been a general trend for diet breadth to be reduced over time (caveats in supplemental text 2).

**Figure 1.**
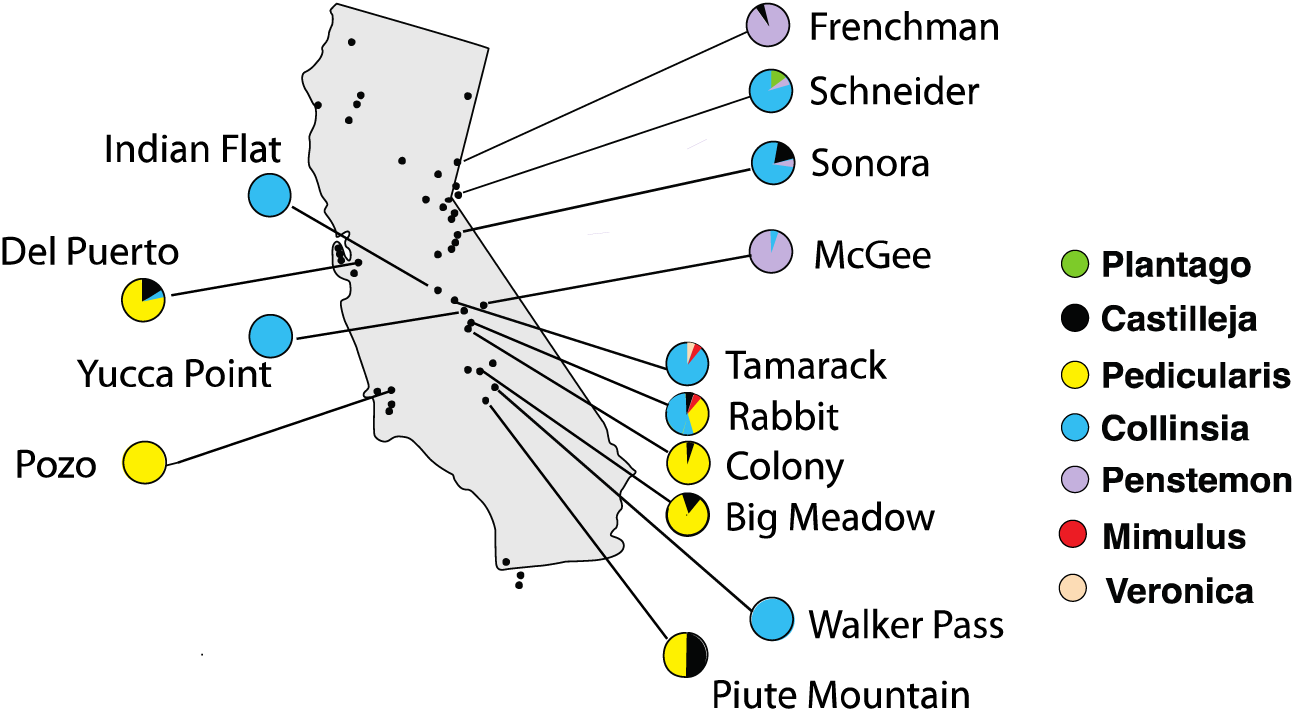
Map of study sites, with diets in 1980’s.

**Figure 2.**
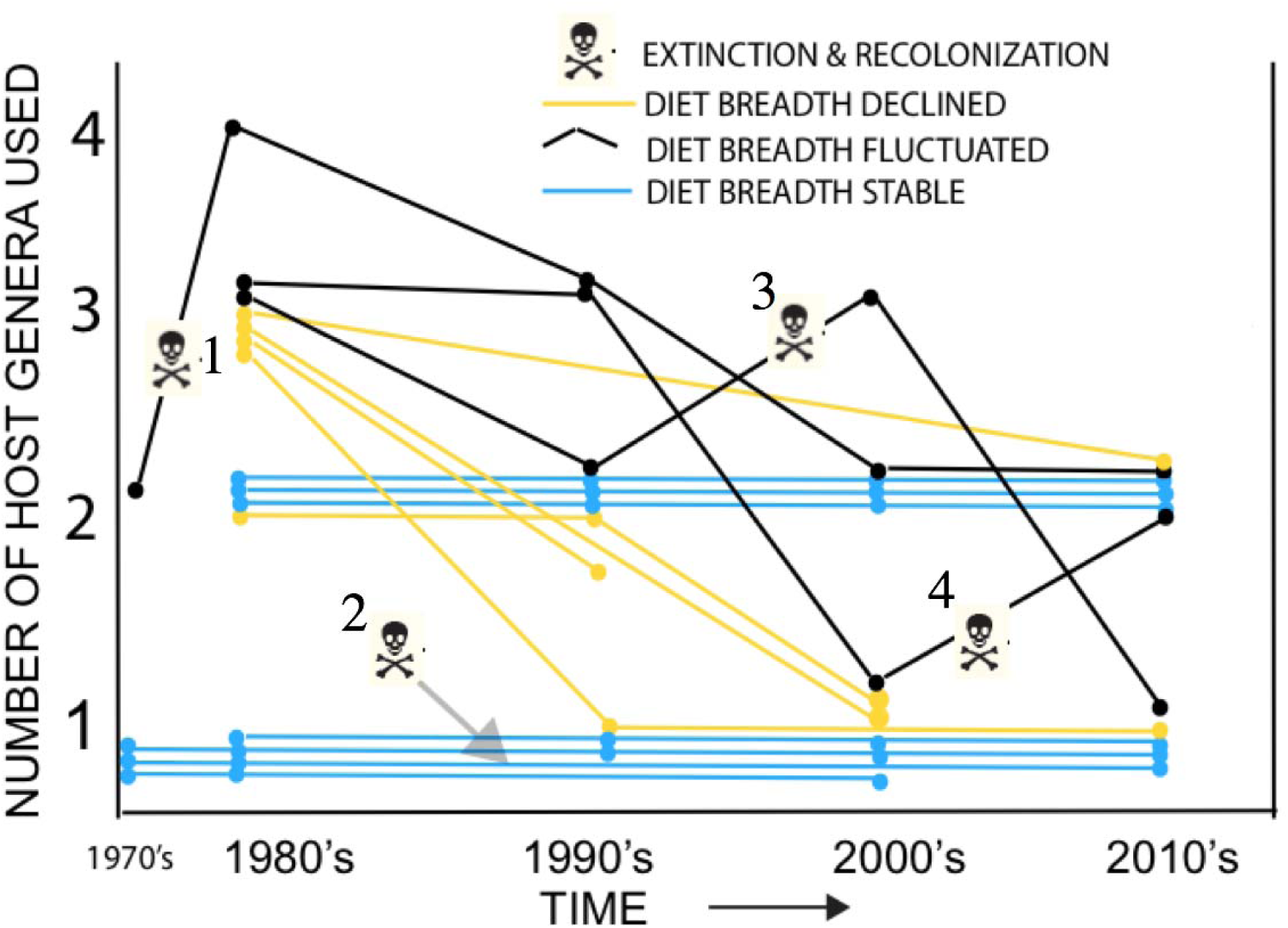
Changes of *E. editha* diet breadth across decades. Extinction/recolonization events numbered by each skull and crossbones: 1 = Rabbit Meadow, 2 = Walker Pass, 3 = Sonora Junction, 4 = Schneider’s Meadow.

### Diet breadth and preference diversity following colonizations

Figure 2 shows extinction/recolonization events at four sites. At Walker Pass only one host was present and the insects unsurprisingly remained monophagous after recolonization. At Schneider a detailed history of diet evolution from 1982-2007 is published^26^ but the apparent diet expansion from one host genus in the 2000s to two after recolonization in the 2010s (Figure 2) depends on a single oviposition^26^ so we shall not discuss it further. At the remaining two sites, Sonora and Rabbit, extinction/recolonization events were clearly followed by broadening of population-level diet driven mechanistically by temporary diversification of oviposition preference ranks. We describe these events in detail below.

#### Sonora: extinction/colonization, diversification of preference ranks and expansion of diet breadth

At Sonora in the 1980’s *E. editha* used three hosts, two of them frequently (*Castilleja, Collinsia*) and one (*Penstemon*) rarely in the 1980s but not at all in the 1990s (Black line number 3 in Figure 2). Host preference ranks were almost invariant; we found a single exception to the rule that butterflies either ranked *Castilleja>Collinsia>Penstemon* or they showed no preference^22,23^. In 1986 the top-ranked host, *Castilleja*, received an estimated 24% of the eggs laid, with *Collinsia* receiving 76%^21^. *Castilleja* was sufficiently rare that most insects failed to find it before reaching the oviposition motivation at which they would accept either *Castilleja* or *Collinsia*, whichever they encountered next. They were then more likely to encounter the more abundant host, *Collinsia*. The principal mechanism by which the population used more than one host was the combination of weakness of preference with rarity of the most-preferred plant^21^.

A natural extinction in the late 1990s, was followed by natural recolonization in 2000-2001, after which preference ranks in 2002 were suddenly diverse (Table 1, Figure 3). We found all possible rank orders for the three hosts, any one of which could be ranked at either the top or the bottom of the preference hierarchy. As expected from these preferences, population-level diet breadth had increased: all three hosts were substantially used in 2002 and the most-used host was *Penstemon*, which had previously been both the least-used and least-preferred of the three (Supplemental text 3).

**Table 1:** Preference ranks at Sonora Junction before and after natural extinction and recolonization.

|  |  | <<<<Prefer<br>plant named<br>at left | No<br>preference | Prefer>>>><br>plant named<br>at right |  |
| --- | --- | --- | --- | --- | --- |
| 1986-88 | Castilleja | 20 | 2 | 0 | Penstemon |
| 1986-88 | Castilleja | 13 | 9 | 0 | Collinsia |
| 1986-88 | Collinsia | 43 | 3 | 1 | Penstemon |
| **Extinction & Recolonization** |  |  |  |  |  |
| 2002 | Castilleja | 7 | 2 | 6 | Penstemon |
| 2002 | Castilleja | 5 | 5 | 5 | Collinsia |
| 2002 | Collinsia | 12 | 2 | 10 | Penstemon |
| 2014 | Castilleja | 21 | 0 | 0 | Penstemon |
| 2014 | Castilleja | 21 | 0 | 0 | Collinsia |
| 2014 | Collinsia | 13 | 5 | 0 | Penstemon |
| 2018 | Castilleja | 28 | 1 | 0 | Penstemon |
| 2018 | Castilleja | 29 | 1 | 0 | Collinsia |
| 2018 | Collinsia | 18 | 5 | 2 | Penstemon |

**Figure 3.**
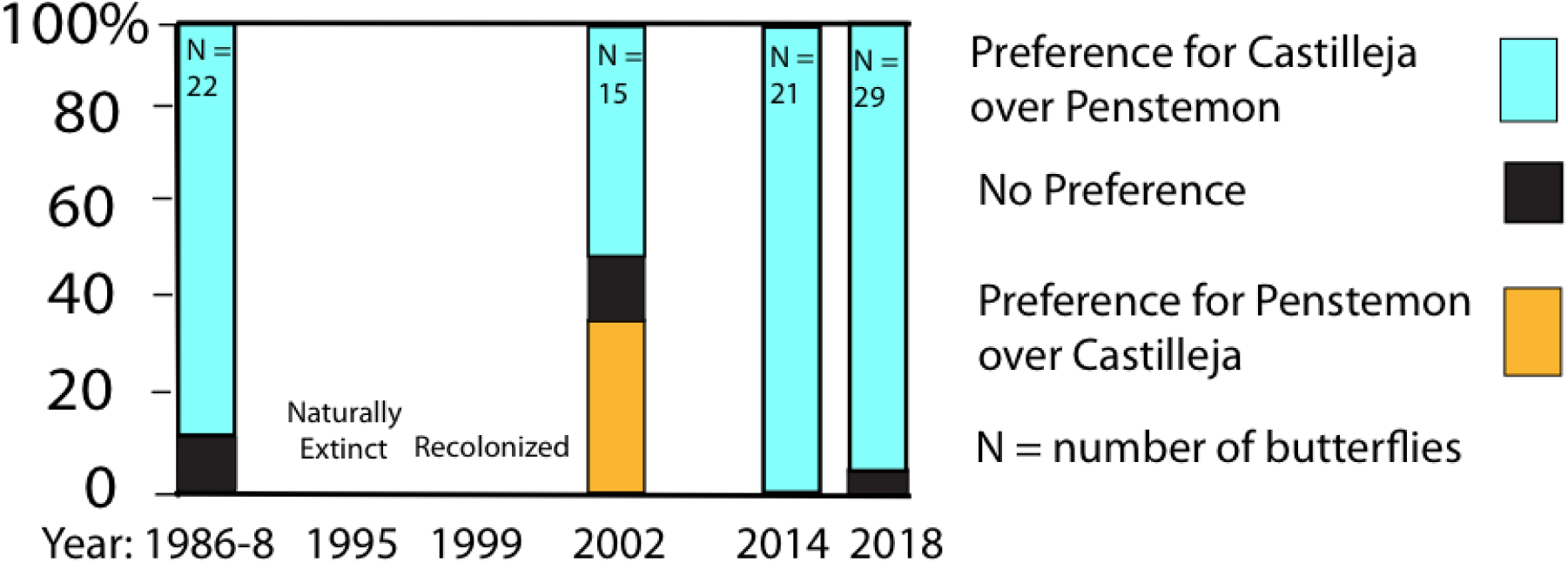
Proportions of butterflies preferring *Castilleja* or *Penstemon* at Sonora before and after natural extinction and recolonization. (additional data in Table 2).

By 2014/2018, with the exception of two butterflies that preferred *Collinsia* over *Penstemon*, preferences at Sonora had reverted to their original homogeneous ranking of *Castilleja>Collinsia>Penstemon* (Table 1, Figure 3). *Penstemon* had once again disappeared from the diet; despite intensive censuses, in neither 2014 nor 2018 did we find a single oviposition on this host. This was not surprising given that, out of 50 females tested, none preferred *Penstemon* over *Castilleja* and only one failed to discriminate between these hosts. Both the diversification of preferences and the inclusion of *Penstemon* into the diet as a major host had been ephemeral, appearing rapidly following the recolonization event, then disappearing within a dozen generations.

Data from 1986-8 are from references 21 and 23; data from 2002, 2014 & 2018 are unpublished.

#### Site: Rabbit: two episodes of diversification of diet and preference: one adaptive, the other nonadaptive

Prior to human intervention, the *E. editha* metapopulation in the Rabbit metapopulation (Fig. 1) used two perennial hosts and occupied >20 habitat patches distributed across 8 × 10 km^27^. The principal diet was *Pedicularis*, with minor use of the much rarer *Castilleja*^28^. A third potential host, the annual *Collinsia*, was abundant but not used. Natural selection opposed using *Collinsia* because its lifespan was so short at this site that larvae hatching from eggs laid on it were almost certain to starve after host senescence^18^.

Between 1967 and 1978, humans made 18 clearings in which all trees were removed, fires were set and ground was bulldozed, locally extirpating the butterflies from the cleared areas. Fertilization effect from the fires extended the size and lifespan of *Collinsia* to the point where they could accommodate the life cycle of the butterflies. *Collinsia* in clearings suddenly became a benign environment for the larvae, supporting higher fitness than the well-defended *Pedicularis*, despite the butterflies being adapted to *Pedicularis* and demonstrably maladapted to *Collinsia* in a suite of six host-adaptive traits^28^.

By the mid-1980s all the larger clearings had been colonized by butterflies immigrating from adjacent unlogged patches, where the insects had persisted on their original diet of *Pedicularis*. In one such clearing a detailed census and map was made of the distribution and host affiliations of *E. editha* oviposition^20^. Eggs had been laid on four hosts: two novel hosts, *Collinsia* and *Mimulus*, plus the two traditional hosts, *Pedicularis* and *Castilleja. Pedicularis* is a hemiparasite of gymnosperms, killed by logging, so just a few individuals had entered the clearing at its margins. *Collinsia* and *Mimulus* were used in the centre of the clearing but remained unused in the adjacent unlogged patch, where both occurred and *Collinsia* was abundant. This pattern of host use sets the context for the two cases of preference diversification that occurred in the clearing and that are detailed below.

Case 1: adaptive diversification of preference as part of host shift from *Pedicularis* to *Collinsia*

Butterflies in Sequoia National Park (12 km from Rabbit) represent the putative pre-logging state of the Rabbit population. Here, we found no diversity of preference rank; most butterflies from the Park showed varying strengths of preference for *Pedicularis* over *Collinsia* and a few showed no preference, but none preferred *Collinsia* over *Pedicularis*^27^.

In contrast, preference ranks for the same two hosts in the anthropogenically altered Rabbit clearing were diverse and evolving through the 1980s. In the early 1980s most insects emerging in the centre of the clearing preferred to oviposit on *Pedicularis*, despite having developed on *Collinsia* from eggs naturally laid on it. The proportion of these butterflies that preferred *Collinsia* increased significantly between 1984 and 1989^27^.

This change, and the diversification of preference from the starting condition lacking diversity of preference rank was measured in the field but also reflected in laboratory-raised butterflies. It is consistent with adaptive evolutionary response to measured natural selection that favoured preference for *Collinsia*, but acted on an initially *Pedicularis-*preferring population^28^.

Case 2: non-adaptive preference diversification: incorporation of *Mimulus* into the diet as a side-effect of host shift to *Collinsia*.

In the ancestral state *Mimulus* and *Collinsia* were present but neither was used for oviposition, though *Collinsia* was fed upon by wandering late-instar larvae. By the early 1980s both plants were hosts in the clearing^20^ (Table 2A) and oviposition preferences for them were diverse (Table 2B). Field experiments^22^ estimated that selection in the clearing favoured oviposition on *Collinsia* over that on *Mimulus*, so *Mimulus* had been included in the diet in the absence of natural selection favouring this addition (*Mimulus* is “host 4” in Fig 2 of reference 22). By the late 1980s preferences for *Collinsia* over *Mimulus* had become homogeneous and *Mimulus* was no longer used (Tables 2A, B). These preferences were still homogeneous in butterflies sampled from undisturbed patches in 2019, 17 years after both *Collinsia* and the clearing patches had been abandoned and the insects had reverted to their traditional diet of *Pedicularis* in the un-cleared patches^28^.

**Table 2A:**
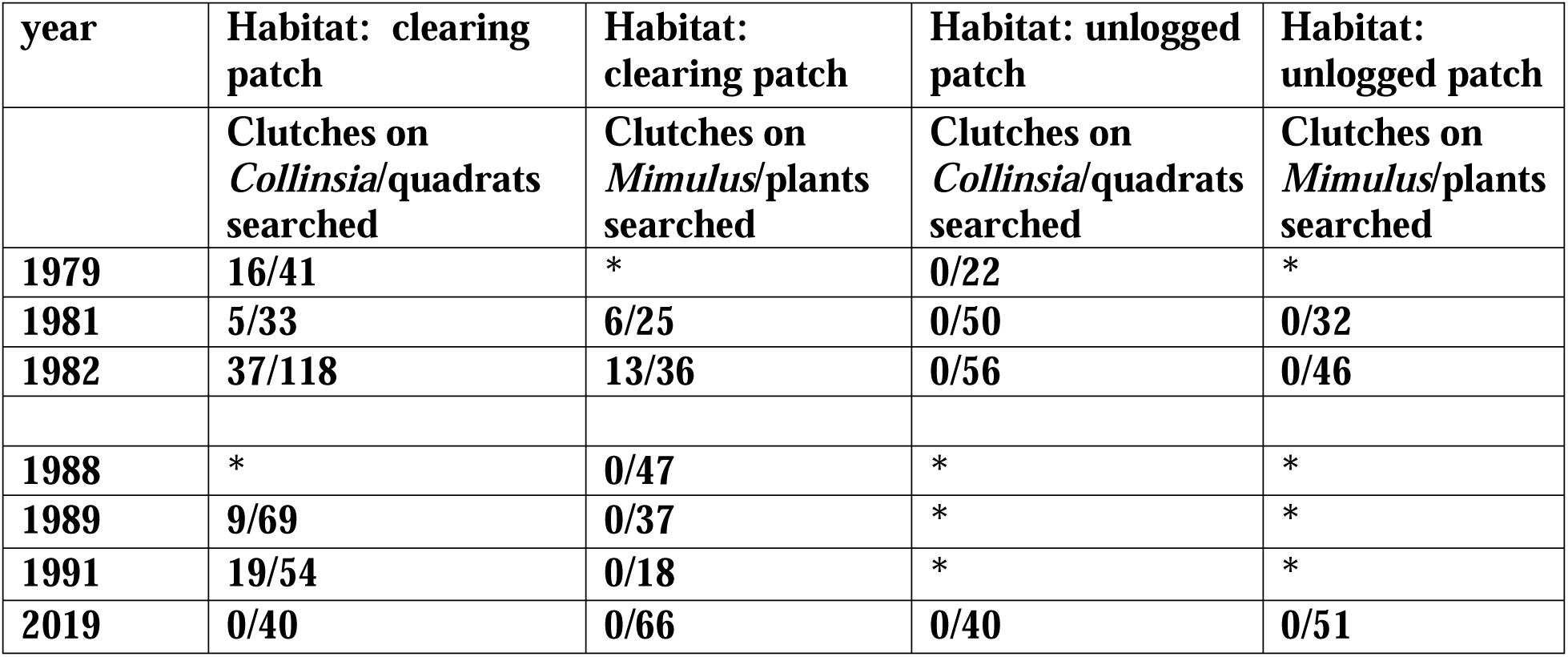
Egg distributions on *Collinsia* and *Mimulus* in Rabbit clearing and adjacent unlogged patch. * indicates that no census was done

**Table 2B:**
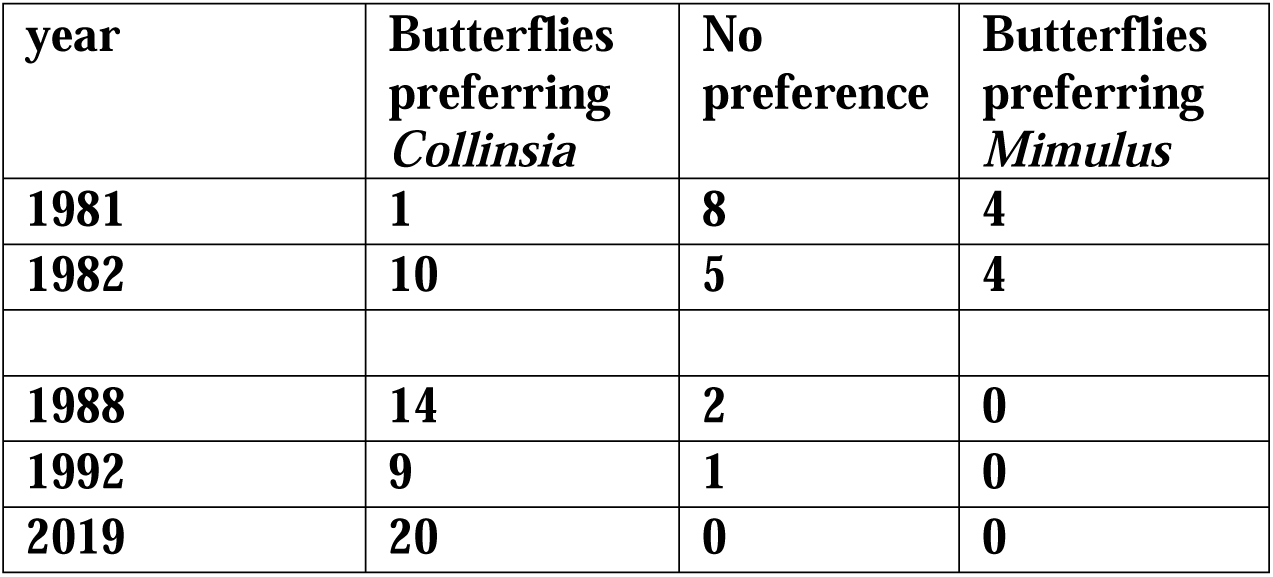
Preferences for *Collinsia* vs *Mimulus* in Rabbit Meadow clearing after its colonization in the 1970’s.

#### Site: Schneider

The traditional diet of *E. editha* at Schneider (Carson City, Nevada, Fig 1) can be deduced from that of the closest known population of the same ecotype, at Simee Dimeh Summit. After fire had caused a brief population explosion of *E. editha* at Simee Dimeh in 2013, we recorded >100 egg clusters on *Collinsia* and none on the only other potential host present, *Penstemon*, despite intensive search. Therefore, our estimate of the starting condition at Schneider is monophagy on *Collinsia*. The same conclusion was drawn by Thomas et al^29^ from the diet of the closest population to Schneider that was then known, at Curtz Lake.

By 1969 the European exotic *Plantago lanceolata* had already been incorporated into the diet alongside *Collinsi*a^26^. We began detailed censuses in the early 1980’s, when three hosts were used: *Collinsia, Plantago* and *Penstemon*. Oviposition preferences were diverse, heritable, and expressed in nature: butterflies captured in the act of oviposition tended to prefer, in preference trials, the host species they had chosen naturally^30^.

From 1982 onwards the population evolved from majority use of *Collinsia* towards increasing use of *Plantago*. By 2005-7 oviposition preferences were invariant, with butterflies unanimously preferring *Plantago* over *Collinsia*. Both *Collinsia* and *Penstemon* had been abandoned. In 2008, the population became extinct in response to a change in land management. In 2013-4 the site was recolonized by *E. editha* adapted to *Collinsia*, on which the new population was initially monophagous^26^.

How does this history fit our scenarios? As at Rabbit, preference tests performed on butterflies in the presumed ancestral condition at Curtz Lake found no diversity of preference rank; butterflies either preferred *Collinsia* over *Plantago* or showed no preference^32,33^. We attribute to natural selection the appearance of preference rank diversity for these two hosts at Schneider in the 1980s and the disappearance of *Collinsia* preference in 2005-7 as the population became monophagous on *Plantago*^26^. It’s possible that *Penstemon* was temporarily included in the diet as a nonadaptive side-effect of the host shift to *Plantago*, in a manner similar to the temporary inclusion of *Mimulus* at Rabbit Meadow.

After recolonization in 2013-4 the population reverted to monophagy on *Collinsia*, failing to exhibit our expected diet breadth expansion^26^. In 2018 we found only two ovipositions, which were on different hosts (Table 1) but we draw no conclusion from this sample.

### mtDNA and diet breadth

In the 1980s, while diet breadth censuses were being done, a separate set of samples was taken for mtDNA analysis, independently of the diet censuses, from 24 (meta)populations of *E. editha*^31^. We here address a question that the original study^31^ did not ask. We test the null hypothesis of no association between genetic diversity and diet breadth. To do this, we need to allow for variation of sample sizes in the genetic study. Accordingly, we derived a sample-size-independent estimate of mtDNA diversity as the number of haplotypes per individual sampled (Supplemental Table 1, right hand column). The association between this statistic and the diet breadths illustrated in Figure 2 and listed in column 2 of Supplemental Table 1 is significant with p = 0.024, by Spearman rank test (two-tailed). As Figure 4 suggests, samples from populations using fewer host genera contained significantly more mtDNA haplotypes.

**Figure 4.**
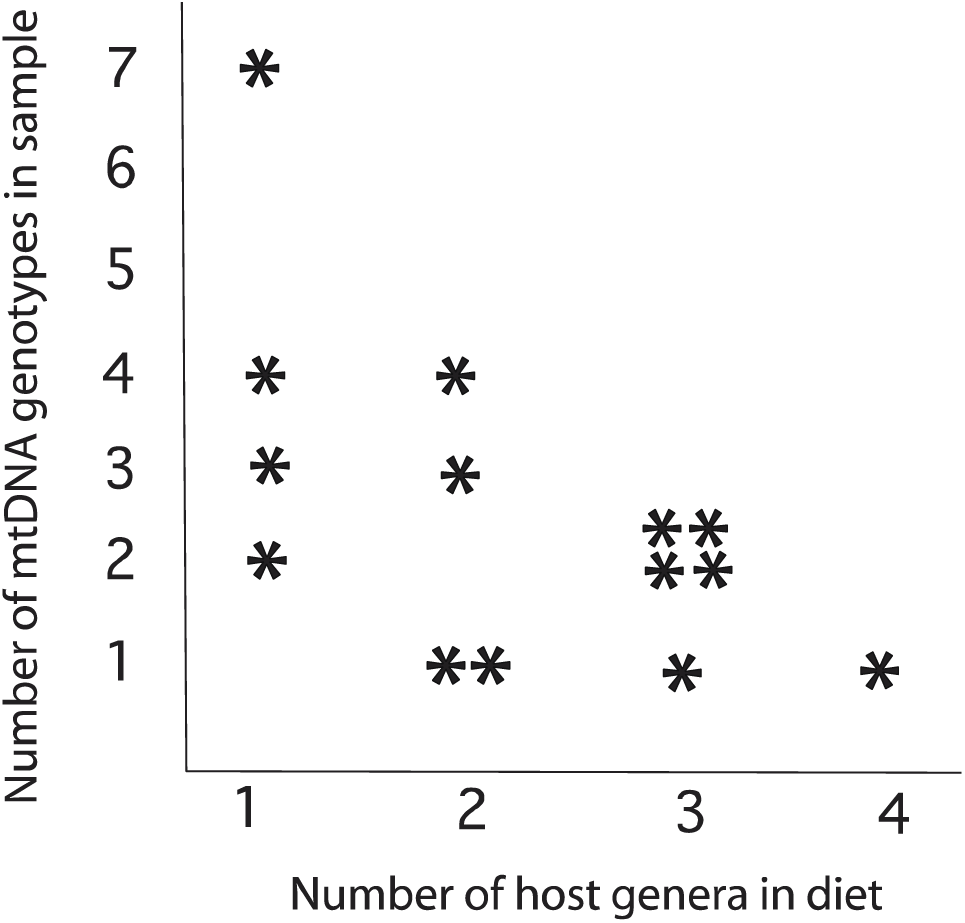
Numbers of mtDNA haplotypes found in the 14 study populations of *E. editha* plotted against the 1980’s diet breadths shown in Figures 1 & 2 and supplemental Table 1.

Because sample sizes were diverse, the association shown in Figure 4 might have stemmed from sampling more individuals from populations that happened to be monophagous than from those with broader diets. However, the opposite was the case: a regression of mtDNA sample sizes on diet breadth, using the data in Table 1, gives a slope of +3.6 (P = 0.06, two-tailed). The direction of this trend, with higher mtDNA sample sizes from populations with broader diets, is opposite to that expected to produce the relationship in Figure 4.

## Discussion

Much of the literature that ties insect diet evolution to generation of biodiversity carries the assumption that host shifts facilitate speciation. In Melitaeine butterflies this does not seem to be true. Host shifts are frequent, closely-related sympatric species typically have overlapping diets^31^, and *E. editha* itself shows strong isolation by distance but no residual isolation by host^19^. The failure of Melitaeines to speciate with host shifts may reflect the fact that they don’t mate on their hosts. Apart from this trait, we have no reason to think that diet evolution in Melitaeines is unusual, so we expect its mechanisms, as revealed in the current study, to be informative about processes that operate more widely than in this butterfly subfamily. Whether the short-term changes we show are informative about long-term diet breadth oscillations that occur across millennia^5-12^ is an open question, but the fact that insects tend to recolonize long-lost ancestral diets suggests that processes measured on very different time scales are related.

### Causes of preference diversification

The first cause of preference diversity that we show is natural selection driving a shift from monophagy on one host towards monophagy on another. We note two previously-published examples from our own work: the generation, during independent host shifts at Rabbit and Schneider, of novel diversity of preference ranks from starting points lacking such diversity. At Schneider we saw this preference diversity appear, persist for >8 years, and then disappear as the insects evolved to monophagy on their novel host^26^. At Rabbit the host shift did not proceed this far. Prior to the metapopulation achieving monophagy on the novel host, the direction of natural selection, and hence the direction of evolution, was reversed, the butterfly populations in the patches using the novel host were extirpated, and the system reverted to its starting point^28^.

The second cause of preference diversity, as a nonadaptive consequence of colonization events, is less expected and not previously published. Our clearest example is the ephemeral inclusion at Rabbit of the unsuitable novel host, *Mimulus*, immediately following the adoption of the suitable novel host, *Collinsia*. The temporary addition of *Penstemon* to the diet at Schneider, during the host shift from monophagy on *Collinsia* to monohagy on *Plantago*, is likely to reflect the same process.

Our results do not apply universally. Two unrelated studies have shown increased dietary specialization after colonization. First, Hardy et al^34^ use phylogenetic analyses to argue that, in scale insects, diet diversity is positively associated with genetic diversity (the opposite of our own finding), so founder effects associated with colonizations and range expansions have caused population-level diet to become more specialized, not less.

The second study with a result contrary to ours is from a butterfly, the Brown Argus, which, like many other species, is indulging in a poleward range expansion attributed to regional climate warming. This expansion is associated with increasing host specialization in England. Oviposition preferences were more specialized and homogeneous, both within and among populations, in the expanding parts of the range than in long-established populations^35^. In addition, larvae in the expanding regions were physiologically more host-specialized and had lost evolvability, compared to their ancestral populations^36^. These two contrary results give us pause in suggesting the level of generality of our results.

However, we regain some confidence because, if even moderately general, the phenomena we document could underpin the broad geographic pattern of diet breadth with decreasing specialization at higher latitudes^13^. Using a global database of diet observations and geographic distributions of Lepidopteran species, Lancaster^14^ concludes that the principal cause of this latitudinal trend is that range expansions cause loss of population-level specialization rather than that generalists make better colonists. Our studies of *E. editha* support this interpretation and further suggest that population-level generalization in the biogeographic data may often represent diversification of specialists rather than (or in addition to) loss of specialization at the individual level.

### Cause of the negative association between mtDNA and diet breadth

What are possible causes of the negative asssociation that we find between genetic diversity and diet breadth? If gene flow and admixture were the main driver of variable diet breadth, populations with broader diets would be those that had received more diverse gene flow, so they should show more, rather than less, genetic variation – opposite to our findings. On the other hand, the observed relationship could be explained if both were functions of population age: that is, if young populations had both lower genetic diversity and broader diets than long-established populations.

Newly-founded populations are, indeed, expected to have reduced genetic diversity and to acquire more genotypes as they age, from some combination of immigration and mutation^32,33^. However, it is not obvious that recently-colonized populations should have broader diets than the sources from which they were derived. Indeed, the opposite relationship can occur. If diet breadth of a source population reflects diversity of individuals with different host adaptations, we expect “specialization by drift”^34^; founder effects should reduce diet breadths at newly-colonized sites, compared to their sources.

One possible mechanism to generate increase in diet breadth after colonizations would be that colonization events were followed by host shifts, and additional hosts were added to the diet during evolutionary transitions from traditional to novel hosts, as suggested by Hardy^9^. A model of parasite evolution in the context of a fitness landscape with heterogeneous hosts does, indeed, generate this scenario^12^.

Colonizing females are unlikely to actively switch host species with high enough frequency to produce the association in Figure 4. However, if, as we have suggested (above and in supplemental text 1), each host population were effectively unique from the butterflies’ perspective, then adapting to a newly-colonized population of a traditional host species might require sufficient change of host preference that additional host species were temporarily drawn into the diet. In this case, an explanation of the diet breadth/mtDNA relationship on the basis of colonizations, host shifts and population age becomes less unlikely (see supplemental text 1).

### Conclusion

As more and more species track shifting climate spaces driven by current warming trends, the numbers experiencing poleward range expansions will continue to rise. Yet we have little understanding of the behavioural and evolutionary processes accompanying these ecological range expansions. The mechanisms driving diet expansion and contraction that we document here may help us to better understand these underlying dynamics, thereby informing projection models and conservation planning under continued anthropogenic climate change.

## Acknowledgements

Paul Ehrlich introduced MCS to *E. editha* in 1967; The U. S. National Park Service permitted work in Sequoia Natioinal Park; Chris Thomas, Helen Billington and David Ng, together with the authors, gathered data shown in Figure 1. Lesley Lancaster, Chris Thomas and Maud Charlery-Massiere critiqued the MS.

Reprint and permissions information is available at www.nature.com/reprints

Correspondence and requests for materials should be addressed to

## Author Contributions

Both authors participated in field censuses and writing. MCS performed oviposition preference tests and statistical analyses.

## GLOSSARY

Population-level diet breadth: In the studies reported here, the number of host species on which eggs of *E. editha* were laid in a particular population.

### Host use

Again, in the work reported here, the proportion of eggs laid on each host species by an insect or an insect population. In a practical sense, this must most often be measured from the distributions of silken webs spun by young larvae, although groups that do not survive to this stage are missed by this technique (see Methods).

Acceptance: a positive behavioural response by an insect to an encounter with a plant. It is a description of an observable and measurable event. It is not a trait of either plant or insect, since it depends on both insect preference and plant acceptability (see below). It is a trait of the plant-insect interaction^38^.

### Insect preference

the set of likelihoods of accepting particular specified hosts that are encountered. Defined in this way, it is a property of the insect that can vary among individuals (Singer 2000) and can be heritable. *E. editha* first encounters hosts visually, then chemically, then physically, with separate preferences expressed at each stage^31^. Again in *E. editha*, the strength of post-alighting preference for two hosts, say host A and host B, is measured by the length of time that a female will search accepting only host B (if encountered) until, after failing to find host B, she reaches the level of oviposition motivation at which either A or B would be accepted, whichever is next encountered (details and justification in reference 39).

### Plant acceptability

The set of likelihoods that a plant will be accepted by particular specified insects that encounter it. Defined in this way, it is a property of the host that can vary among individuals^38^ and can be heritable^23^.

## Methods

### MtDNA analyses

Reference 26 used 17 restriction endonucleases to identify 22 mtDNA haplotypes of *E. editha*, the distributions of which were recorded within and among 24 populations or metapopulations of the butterfly, with metapopulations treated as single sites. These sites varied dramatically in their haplotype diversity. (Meta)populations with sample sizes of 11, 13, 17 and 30 each contained single haplotypes, while a sample of 14 individuals produced 7 haplotypes and a sample of only four contained no replicates (Table 1). In order to retain this last small but informative sample we chose to include in our analysis all populations with mtDNA sample sizes of four or greater, thereby reducing the number of populations analyzed from 24 to 14.

### Preference tests

Butterflies were captured in the field and their oviposition preferences tested by a standardized technique, in which encounters are staged between the tested insect and each plant in alternation. Plants were undisturbed in their natural habitats or freshly transplanted into pots in their own soil. Acceptance of plant taste was judged from full abdominal curling and extrusion of the ovipositor for 3 sec. Acceptance and rejection were recorded at each encounter, but oviposition was not allowed. Videos showing acceptance in such staged encounters are in Singer & Parmesan 2019. During each test the range of plants that would be accepted, if encountered, expands over time with increasing motivation to oviposit. Therefore, acceptance of plant A followed by rejection of plant B is recorded as preference for A over B. Testing of assumptions underlying this technique described in reference 32. Insects without preference are not shown in the Figure, so percentages do not sum to 100% except in 2005 & 2007, when preference for *Plantago* was unanimous among tested butterflies. Raw data are in Extended data table 3. A more detailed comparison between early and late periods, showing strengths as well as direction of preference, is given in Extended data Fig.8. The assumption that these insects’ preferences are not influenced by prior experience, either as larvae or as adults, is supported by prior observation and experiment^40^.

Supplemental text 1.

### Complex variation of host acceptability from the butterflies’ perspective

1. *M. cinxia* were subjected to staged encounters with hosts during each of which they tasted the plant and could respond by attempting to oviposit (host acceptance) or by basking (host rejection). They were asked to rank three individual *Plantago lanceolata* and three *Veronica spicata* plants. Some individual butterflies consistently ranked these hosts taxonomically, either ranking all three *Plantagos* above all three *Veronicas* or vice versa. Other butterflies from the same populations were more impressed with chemical variation among individual hosts than between species, consistently producing rankings in which the two species were interdigitated^24^.
2. *E. aurinia* produced different rankings of host genera, depending on which individual plants the butterflies were offered. This effect produced the false impression that insects from two monophagous populations, one using *Lonicera* and the other using *Cephalaria*, preferred to oviposit on *Succisa*, a plant that was not present in either of their habitats^25^.
3. The choice by *E. editha* of different host species at two sites with similar vegetation was shown to be driven approximately equally by genetic variation in host acceptability between sites and by genetic variation in butterfly preference^23^.

### Colonization and host shifts

The studies summarized above all show that, from the butterflies’ perspective, within-host-species variation can complement or even overwhelm variation between host genera. With this in mind. suppose that an emigrant from a population monophagous on *Penstemon rydbergii* colonizes a new habitat, where, given her initial host adaptations, the host that supports highest fitness is likewise *P. rydbergii*. However, the *Penstemon* population at the colonized site differs sufficiently in chemistry from the conspecifics at the source site, that the immigrant female perceives it just as differently as she would perceive a different host species. Colonizing what appears to humans as the same host species would be, to the female, colonizing a different host entity, and this difference could drive diversification of preference and temporary taxonomic broadening of diet, pending natural selection causing a return to monophagy on *Penstemo*n. This hypothetical scenario gains credibility since we have shown that heritable interpopulation variation in acceptability of *P. rydbergii* to *E. editha* was sufficient to cause the butterflies to use the *Penstemon* at one site but to drive them onto a different host genus (*Collinsia*) at another, where the *Penstemon* was less acceptable^23^.

### Prior publication

Some of the ideas presented here were foreshadowed in 2008^41^, including a verbal description of the relationship shown in Figure 2; but no data or analyses were provided at the time.

**Supplemental Table 1:**
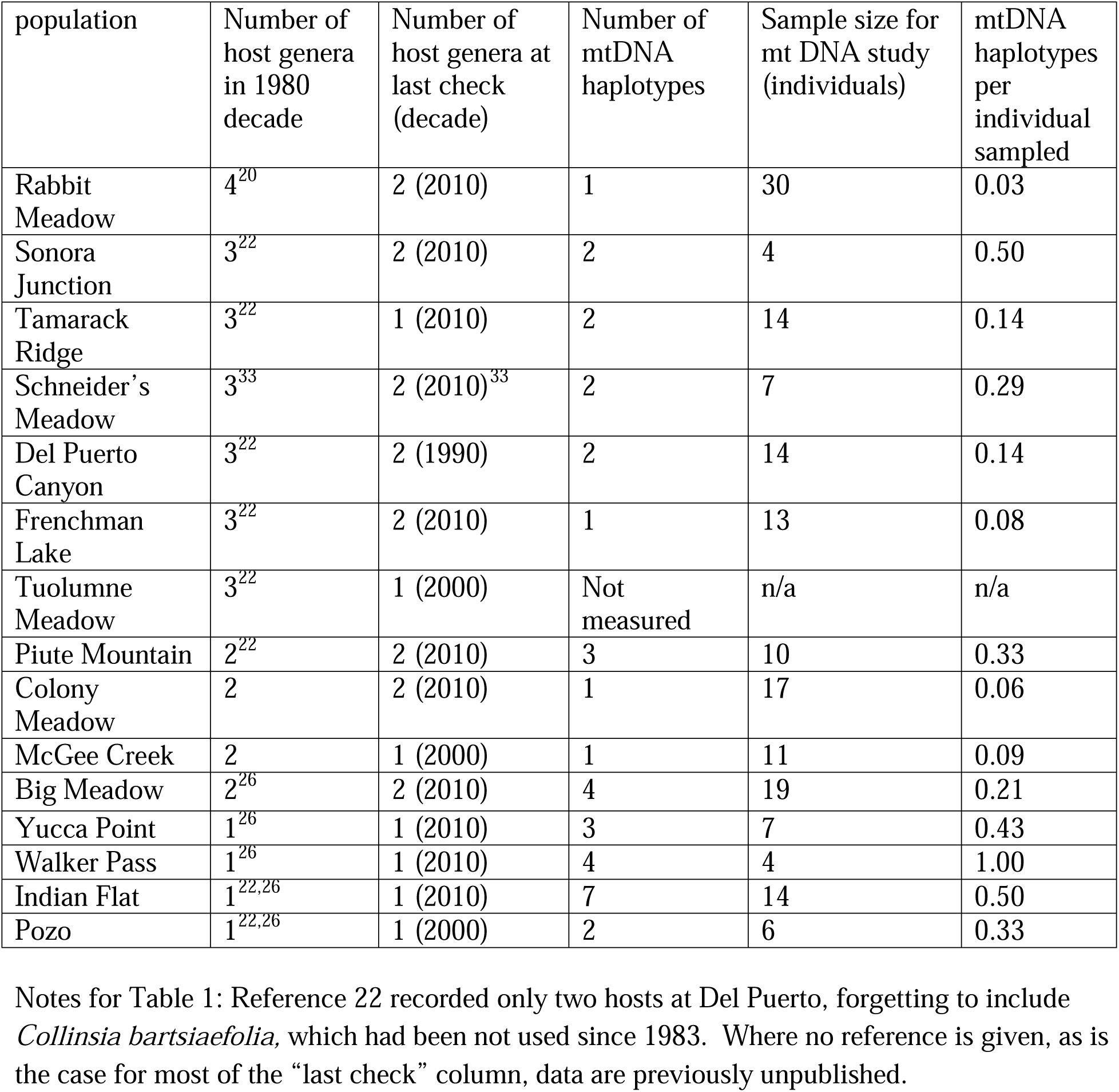
diet breadths in 2 time periods and mtDNA diversity in 1980s.

| population | Number of host genera in 1980 decade | Number of host genera at last check (decade) | Number of mtDNA haplotypes | Sample size for mt DNA study (individuals) | mtDNA haplotypes per individual sampled |
| --- | --- | --- | --- | --- | --- |
| Rabbit Meadow | 4<sup>20</sup> | 2 (2010) | 1 | 30 | 0.03 |
| Sonora Junction | 3<sup>22</sup> | 2 (2010) | 2 | 4 | 0.50 |
| Tamarack Ridge | 3<sup>22</sup> | 1 (2010) | 2 | 14 | 0.14 |
| Schneider's Meadow | 3<sup>33</sup> | 2 (2010)<sup>33</sup> | 2 | 7 | 0.29 |
| Del Puerto Canyon | 3<sup>22</sup> | 2 (1990) | 2 | 14 | 0.14 |
| Frenchman Lake | 3<sup>22</sup> | 2 (2010) | 1 | 13 | 0.08 |
| Tuolumne Meadow | 3<sup>22</sup> | 1 (2000) | Not measured | n/a | n/a |
| Piute Mountain | 2<sup>22</sup> | 2 (2010) | 3 | 10 | 0.33 |
| Colony Meadow | 2 | 2 (2010) | 1 | 17 | 0.06 |
| McGee Creek | 2 | 1 (2000) | 1 | 11 | 0.09 |
| Big Meadow | 2<sup>26</sup> | 2 (2010) | 4 | 19 | 0.21 |
| Yucca Point | 1<sup>26</sup> | 1 (2010) | 3 | 7 | 0.43 |
| Walker Pass | 1<sup>26</sup> | 1 (2010) | 4 | 4 | 1.00 |
| Indian Flat | 1<sup>22,26</sup> | 1 (2010) | 7 | 14 | 0.50 |
| Pozo | 1<sup>22,26</sup> | 1 (2000) | 2 | 6 | 0.33 |

Notes for Table 1: Reference 22 recorded only two hosts at Del Puerto, forgetting to include *Collinsia bartsiaefolia*, which had been not used since 1983. Where no reference is given, as is the case for most of the “last check” column, data are previously unpublished.

The three left-hand columns of the Table show population names, the numbers of host genera on which *E. editha* eggs or larval webs were found during the 1980’s and the numbers of genera used in the most recent decade of observation, which, with four exceptions, is the current decade.

Supplemental text 2

We interpret the trend of decreasing diet breadth over time (Table 1) with caution, for two reasons. The first reason is that diet breadths recorded in the 1980s at two sites, Sonora and Frenchman, depend on single observations of natural egg clutches on the least-used hosts (*Penstemon* at Sonora and *Collinsia* at Frenchman). The second reason is that the broader 1980s diet at one site, Del Puerto Canyon, clearly reflected plasticity of the insects, which temporarily added *Collinsia* to their diet in 1983, when unusually high precipitation rendered this host phenologically suitable by extending its lifespan. *Collinsia* had not been not used at Del Puerto in our earliest observations, in 1969, so its absence from the diet in the most recent observation likely reflects its low acceptability, rather than increased specialization of the insects. Diet breadth at a second site, McGee, has also oscillated during our study, so the low diet breadth in the most recent observation may be coincidental. However, removal of both Del Puerto and McGee from the dataset does not eliminate significance of the trend for recent diets to be more narrow than those originally recorded, within continuously-occupied sites (p = 0.03).

Since the mtDNA data in Figure 2 were compiled from long-outdated techniques, it would seem logical to re-sample the populations with better methodology. However, as Table 1 shows, the diet breadth diversity that existed in the 1980s in our set of study populations has diminished and we longer have the necessary variation of diet breadth among these populations’ to ask the question. If our thesis is correct, some of the new populations founded over the study period should have broader diets, and the mean population-level diet breadth that existed in the 1980s may have been maintained across California. However, to ascertain this we would need to have performed systematic searches for newly-founded populations. We have not done this. Instead, our work has concentrated on the study sites that we first identified between 1968 and 1992.

Together with the mitochondrial data, this evidence suggests that dietary breadth increases after colonization, then declines as a function of population age. These indirect sources of evidence are supported by direct observations of the process, presented below.

Supplemental text 3: At Sonora in 2002 we found 20 egg clutches on *Castilleja* in a total census of this rare plant; 9 on *Collinsia* in a census covering approximately 40% of phenologically-suitable plants and 14 on *Penstemon* in a census covering about 20% of these plants. From these data we estimate that the most-used host was *Penstemon*.

